# Increasing information gain in animal research by improving statistical model accuracy

**DOI:** 10.1101/2022.06.03.494726

**Authors:** Christoph Waterkamp, Vanessa Tabea von Kortzfleisch, Christoph Neu, Simo Kitanovski, S. Helene Richter, Daniel Hoffmann

## Abstract

Reduction of the numbers of laboratory animals is one of the three pillars of ethical animal research. Equivalently, information gain per animal should be maximized. A road towards this goal that is barely taken in current animal research is the more accurate statistical modeling of experiments. Here we show for a typical experiment (“open field test”) with outcomes that are non-normally distributed count data, how this can be implemented and what information gain is achieved. We contrast the state of the art – the use of confidence intervals based on null-hypothesis significance testing (NHST) –, with a Bayesian approach with the same underlying normal model, and a Bayesian approach with a more accurate negative binomial model. We find that the more accurate model leads to a marked improvement of knowledge gained with the experiment, especially for small sample sizes. As experimental data that violate assumptions of simple, conventional models are frequent, our findings have wider implications.

## 1. Ethical concerns in animal research

It lies in the very nature of biology, and in extension, of medicine, that it uses animals as objects of experimental research. However, animals like mice, dogs, or monkeys are not only objects but also subjects, sentient individuals that can suffer from experimental procedures [1]. Therefore, there is a tension between the value of gaining scientific knowledge and avoidance of suffering. To ease this tension, ethical animal research follows the “3R” principle [2]: (1) to *replace* animals as research objects, e.g. by cell cultures or simulation; (2) to *refine* experiments by minimising the pain, suffering, distress or lasting harm that research animals might experience; (3) to *reduce* the number of animals by maximizing the information gain per animal. In the work presented here we will focus on the last aspect, *reduction*. Thus our guiding question is: How can we maximize information gain per animal?

## 2. An example: the open field test

We illustrate our work with a classic behavioral test, the Open Field (OF) test [3], in which individual animals are placed in a small arena and their behavior is monitored (Fig. 1). One possible readout of the experiment is how often the animal explores the center of the arena (“center entries”) in a fixed time interval. An animal that is more anxious may stay close to the walls of the arena and thus has a lower number *y* of center entries. Usually, OF experiments are conducted with differently treated animals to characterize the impact of the treatment on animal behavior, e.g. the effect of chemicals [4], environment [5], or genotype [6].

**Figure 1:**
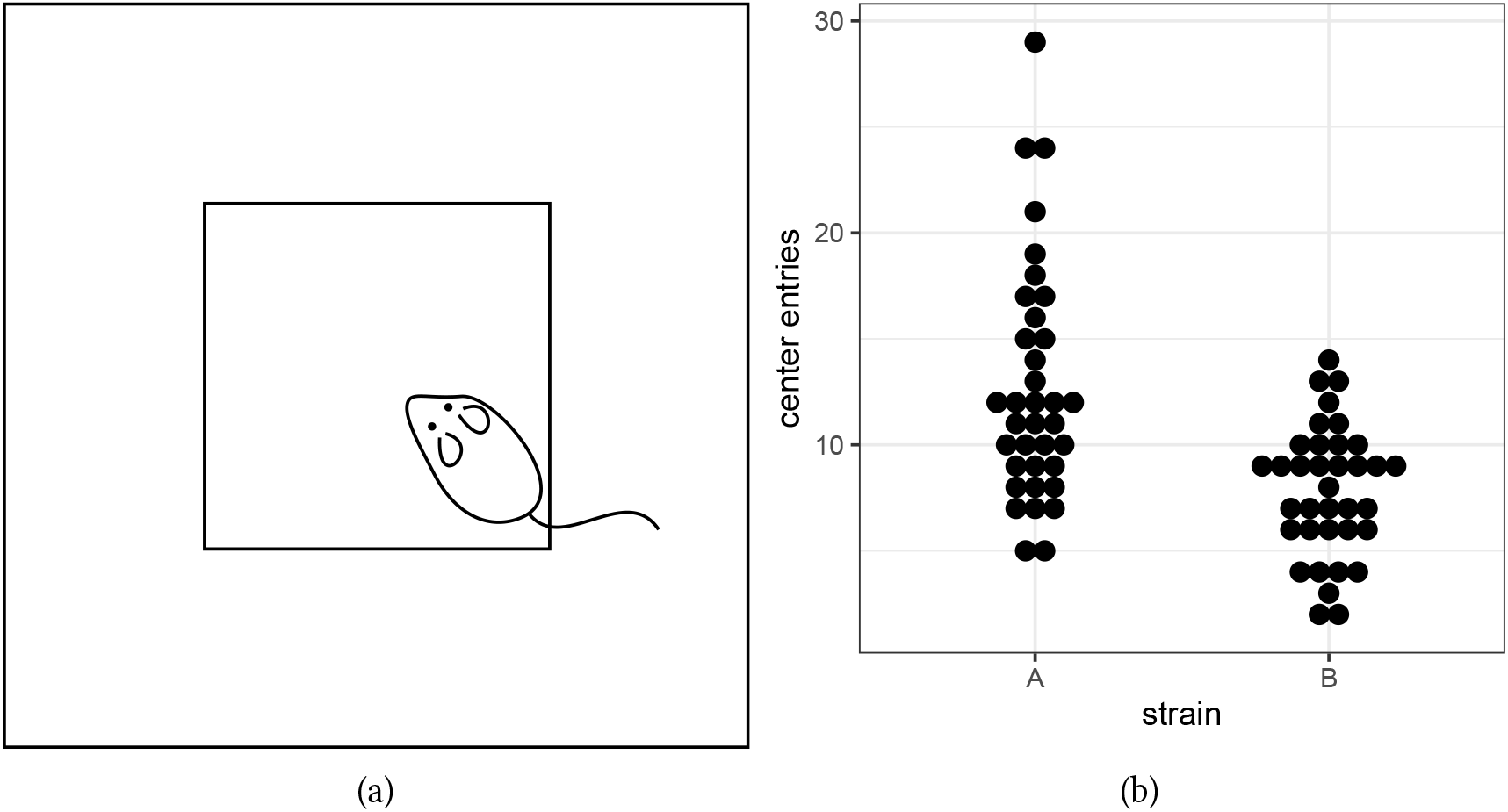
Open Field (OF) experiment. (a) Schematic representation of experimental set-up. Arena and centre areas are indicated by the outer and inner squares, respectively. In this case the mouse has entered the center. (b) Example data from [6]. Each point corresponds to the number of center entries of an individual mouse during a time interval of 5 min. We compare center entry numbers of two genetically different mouse strains, A and B, with each strain represented by 36 animals.

The data in Fig. 1b shows a clear effect of the strain on the number of center entries: individuals of strain A have on average more center entries than individuals of strain B. Empirically, we find with the data in Fig. 1b center entry averages of 〈*y_A_*〉 ≈ 12.7 and 〈*y_B_*〉 ≈ 7.8 for strains A and B, respectively, so that the effect of strain is about *δ* = 〈*y_B_*〉 – 〈*y_A_*〉 ≈ −5. However, we cannot be sure of these estimates as the variation is considerable and there is a large overlap of entry counts between strains. We also observe in Fig. 1b that the distribution of *y_A_* is positively skewed, i.e. we have a heavy upper tail, while *y_B_* could be normally distributed; the empirical values of the skewness are *b*_1*A*_ ≈ 1-0 and *b*_1*B*_ ≈ −0.1, for A and B respectively.^1^ A positive skew is to be expected as *y* are strictly non-negative counts whereas there is no hard upper bound – some mice are apparently very active and reach high center entry counts.

## 3. Current statistical practice in animal research is severely flawed

The structure of the data in Fig. 1b with the differences in distribution variance and shape between *y_A_* and *y_B_* violates assumptions underlying conventional null hypothesis significance testing (NHST) procedures such as t-tests [8] (assumption: normality, homogeneous variance) or Mann-Whitney U test [9] (assumption: same shape and variance of A and B). It is also noteworthy that these tests are of limited value for small sample sizes. However, limitations like these are not the only drawbacks of NHSTs, not even the most severe ones [10]. In particular, the widespread, uncritical and naive application of NHST with p-values in biology and medicine with their noisy data, small effects, and often small sample sizes has contributed to the so-called replication crisis in these disciplines [11], leading to calls to overcome NHST with p-values [12]. In the context of animal research, we note that laboratory animals used in experiments for which results cannot be replicated have been used in vain, which, of course, is the opposite of *reduction* [2].

One of the flaws of NHST with p-values is the “significance” criterion in which the probability *p* of observing the measured effect under the null hypothesis is compared to an arbitrary threshold of 0.05, the famous *p* < 0.05 criterion. With this criterion, measurements on some meaningful scale are artificially dichotomized into the often meaningless classes “significant” and “not significant”.

In basic research, where we are often interested in quantitative knowledge and not in artificial classes, this procedure can be improved by omitting the last step, the application of the “significance” criterion, and instead quantifying likely effect sizes. This emphasis on quantitative estimation lies at the heart of several proposals on how to reform statistical practice [13, 14, 15]. In the following we will therefore rely on methods that estimate effect sizes with their uncertainties.

## 4. Effect size estimation with uncertainty: confidence intervals and highest density intervals

The most commonly used statistical model for life science data is the normal distribution. It underlies the t-test [16,8], ANOVA [17], or the conventional confidence intervals (CIs) as favored by [13]. We follow the latter reference and compute CIs as they emphasize the uncertainty of results and are mapped on the biologically meaningful scale of effect sizes.

Specifically, we calculate 95% CIs of the effect *δ* = 〈*y_B_*〉 – 〈*y_A_*〉 of the mouse strain on the center entries count *y*. The 95% CI is the interval of *δ* estimates for which *p* ≥ 0.05, with *p* the p-value computed for a two-sided Welch t-test[8]. For instance, if *δ* = 0 lies in the 95% CI, we cannot reject the hypothesis that the effect of the mouse strain on center entries count *y* is zero.

One drawback of CIs is that they do not differentiate between points within the CI. Intuitively we expect that *δ* values at the boundaries have a lower probability than *δ* values in the middle, but the underlying NHST makes no statement about the probability of hypothetical *δ* values [18]. An alternative method that allows this is Bayesian inference (for introductory texts see e.g. [19, 20]). Bayesian inference produces so-called posterior probabilities, i.e. conditional probabilities *p*(*δ|y*) of parameter values (here: *δ*), given the measured data *y*. We compute 95% highest density intervals (HDIs), i.e. intervals on the *δ* scale for which the probability *p*(*δ|y*) is greater than for any *δ* value outside the interval.

Despite the fundamental conceptual differences between CIs and posterior HDIs [18] the locations and widths are similar in this case (Fig. 2). Both 95% CI and 95% HDI of the normal model lie around about *δ* ≈ −5 and clearly exclude *δ* = 0, which was to be expected as the effect of the strain on center entries *y* was obvious in the data (Fig. 1b).

**Figure 2:**
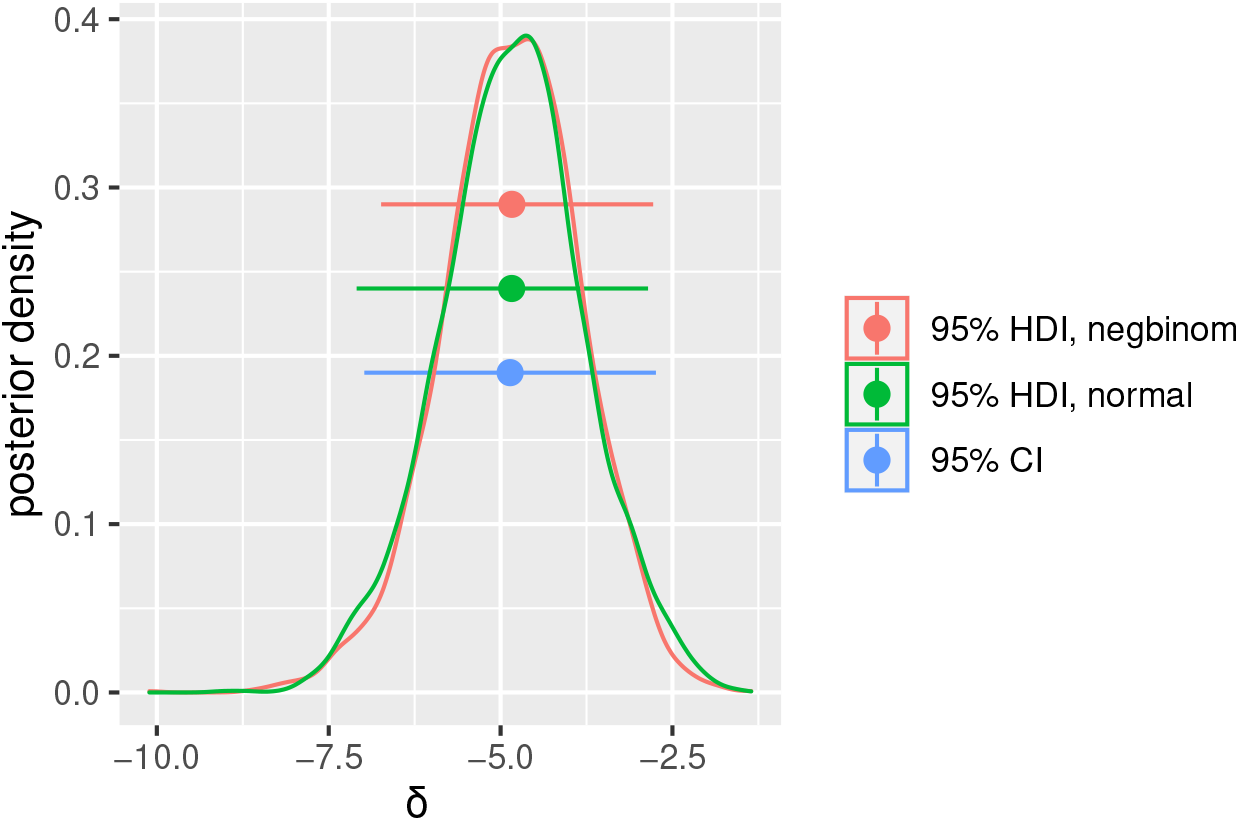
Uncertainty intervals and probability density of strain effect size *δ*. Horizontal bars are 95% HDI of negative binomial model (red bar), 95% HDI for normal model (green bar), and 95% CI (blue) also based on a normality assumption. The two curves represent Bayesian posteriors of the negative binomial model (red curve) and the normal model (green curve). Intervals and densities have been computed for the same OF center entries data in Fig. 1 b.

What we did technically to produce Fig. 2 was to extract 95% CIs from the output of Welch’s t-test [8] in R [21]. For the Bayesian results we formulated the model in the probabilistic programming language Stan and numerically computed the posterior distribution by Hamiltonian Markov Chain Monte Carlo sampling [22] (3 chains, each with 1000 steps of warm-up and 1000 steps of sampling). A weakly informative prior was used, i.e. a prior that allows exploration of posterior regions somewhat larger than is realistic, as was confirmed by prior predictive checks [23]. New data was simulated from the posterior and found for the normal model in reasonable agreement with observed data (posterior predictive check, Fig. 3a). The Stan program for the normal model is given in Appendix A.

**Figure 3:**
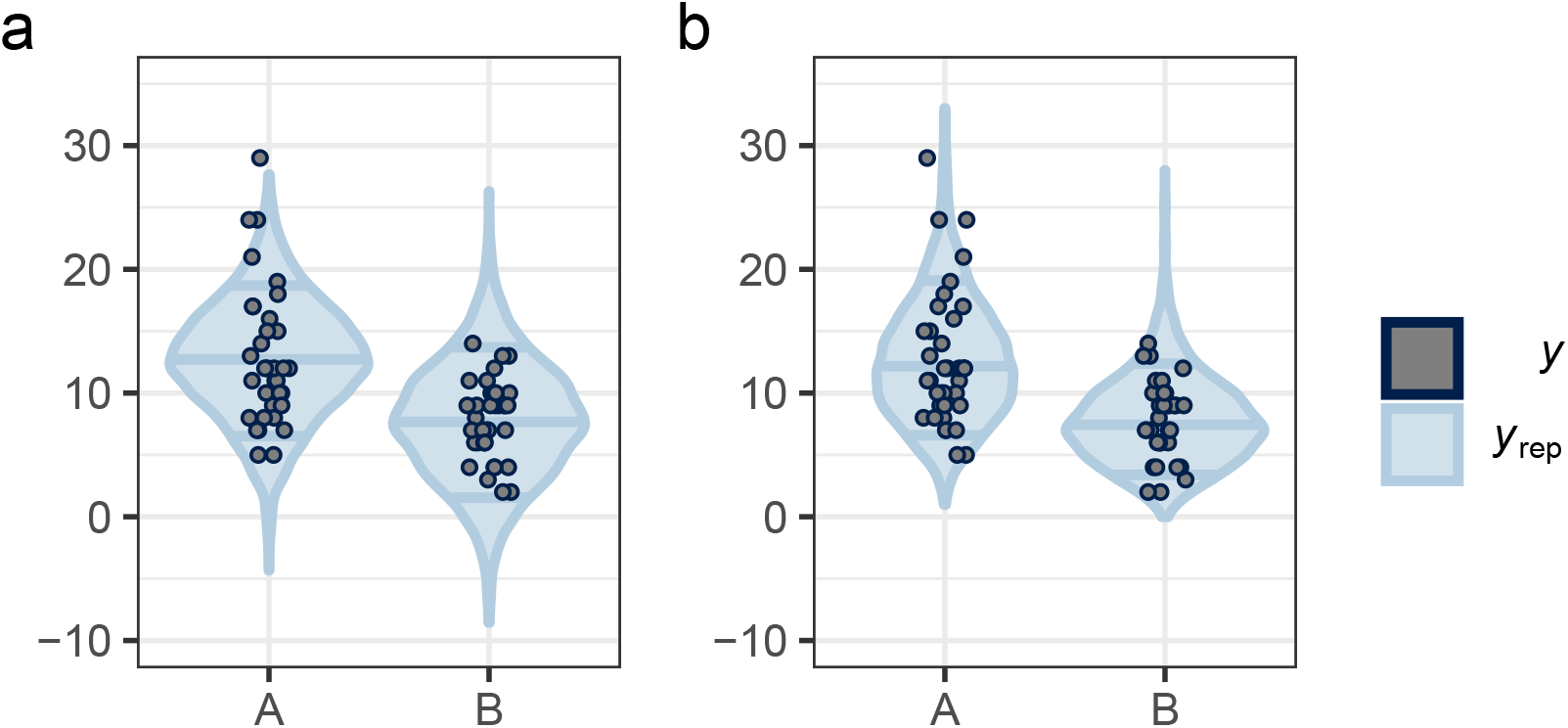
Posterior predictive checks for the normal model (a), and the negative binomial model (b). Counts of center entries (vertical axis), simulated with the Bayesian models for both mouse strains, A and B (horizontal axis), are compared with observed counts from Fig. 1 b. Dark points: observed counts *y* (with random jitter of data points along the horizontal axis to reduce overplotting); blue violins: probability densities of the simulated data *y*_rep_. Horizontal blue lines in the violins mark 0.25, 0.5, and 0.75 quantiles of the replicated data.

## 5. Mapping the experiment onto a more accurate model

A close inspection of Fig. 3a shows some shortcomings of the normal model. The model assumes normally distributed *y* values. This means that it predicts density on all of [−∞, ∞]. In fact, we see in Fig. 3a probability density for *y* < 0 for both strains, A and B. However, we know that *y* are counts, i.e. *y* ∈ ℕ_0_. Conversely, the normal model misses the positive skew in the data described in section 2. Thus, the normal model predicts very low density in a region where we see data points, especially at the highest count in strain A, *y_A_* = 29.

A simple correction would be to model the data by two Poisson distributions 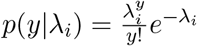 with *i* ∈ {*A, B*}. This would fix qualitatively the issues of the *y* range and skew. This would work for simple physical objects with strain as the only degree of freedom but not for biological organisms with their complex, variable behavior. Thus, for each strain we effectively have not one Poisson distribution with a single parameter λ but a mixture of Poisson distributions with different λ values. Often, it is assumed that λ follows a Gamma distribution, so that we have a so-called Gamma-Poisson mixture distribution of counts for each strain [20], or, as it is more widely known, a negative binomial distribution:

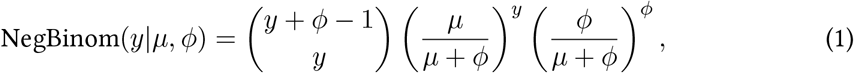

with parameters *μ* ∈ ℝ^+^ and *ϕ* ∈ ℝ^+^ we have expectation value 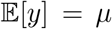 and variance 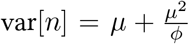. Note that for *μ*^2^ ≪ *φ* both variance and mean become *μ*, i.e. we recover the features of a Poisson distribution; the precision parameter *φ* tunes over-dispersion. It is easy to formulate a Bayesian model with likelihood Eq 1 (Stan program in Appendix B), to run Bayesian inference and simulate new data as described before for the normal model. The posterior predictive check in Fig. 3b shows a better fit between measured and simulated data. The density of *y*_rep_ reproduces the positive skew seen in the measurements and of course, does not predict negative counts, in contrast to the normal model.

There are, however, only small differences between the 95% CI, the 95% HDI from the normal model, and the 95% HDI from the negative binomial model (red in Fig. 2). It appears that the 95% HDI from the negative binomial model is a bit narrower and the probability density of *δ* a bit more concentrated around the mean *δ* with thinner tails to both sides than the normal density. So, we see in the plots only a weak effect of the model on the inferred effect size *δ*, and it appears that little more information about the effect size can be gained by using the more accurate model.

A way of assessing model performance is to quantify its ability to generalize to unseen data. Given that the normal model predicts density of *y* in nonsensical regions and ignores *y* regions that matter, we should expect that the negative binomial model generalizes better. This is in fact what we find in a direct comparison of the two Bayesian models by leave-one-out cross-validation [24]: The expected log pointwise predictive density is by 8.7 lower for the normal than for the negative binomial model, with a standard error of 3.2, i.e. the negative binomial model generalizes better than the normal one.

A critical factor that we have neglected so far is the sample size. A sample size of 36 animals per group is quite substantial, given that the effect size is considerable. It would be very surprising if under these conditions several reasonable methods would not come to similar results. However, in published biological experiments, sample sizes of 3 to 6 animals per group are not unusual. Thus, if we are interested in *reduction*, we must compare the three methods or models for smaller sample sizes.

## 6. Numerical experiments to quantify the impact of model accuracy on information gain for small sample sizes

Conceptually, NHST and Bayesian inference are fundamentally different and produce different kinds of outcomes, which makes direct comparisons difficult. However, in both paradigms we can express likely outcome values and their uncertainties by intervals, e.g. 95% CIs from NHST and 95% HDIs from Bayesian inference [13, 18]. Thus, we can compare the performance of methods and models by comparing these “uncertainty intervals”. In this section, we present such a systematic comparative analysis for the small sample sizes that are typically used in animal research.

For the OF test data in Fig. 1b, we have shown that this type of data can be modelled by a negative binomial distribution. With our Bayesian procedure introduced previously, we found for the parameters in Eq 1 values of *μ_A_* ≈ 12, *μ_B_* ≈ 7, and *ϕ* ≈ 10 (assuming that there is only one value *ϕ* for both mouse strains A and B). We used these realistic parameter values to simulate random sets of *y_A_* and *y_B_* values of the corresponding two negative binomial distributions with sample sizes of *n* = 3, 5, 7,…, 31 animals per strain. For each of these 15 sample sizes *n* and both strains A and B, we generated 1000 of these data sets *y_Ani_, y_Bni_* with *i* ∈ {1,2,…, 1000}. We then computed uncertainty intervals for each pair of sets *y_Ani_,y_Bni_*, i.e. 95% CIs (from t-tests) and 95% HDIs (from Bayesian inference with normal and negative binomial models) as described before. Finally, we compared the resulting uncertainty intervals, the distributions of their widths and lower and upper bounds.

### 6.1. Better model accuracy leads to narrower uncertainty intervals

We found that for the smallest sample sizes, in particular for *n* = 3, which is frequently employed in biomedical research, there is a strong influence of the three methods on the distributions of lower and upper bounds (Fig. 4). With increasing sample size *n* per mouse strain, the distributions of lower and upper bounds become quickly more narrow and also more similar across the three compared methods. For the 95% HDI of the negative binomial model, the distributions of lower and upper bounds are consistently most narrow and also closest to the true value of *δ* = −5.

**Figure 4:**
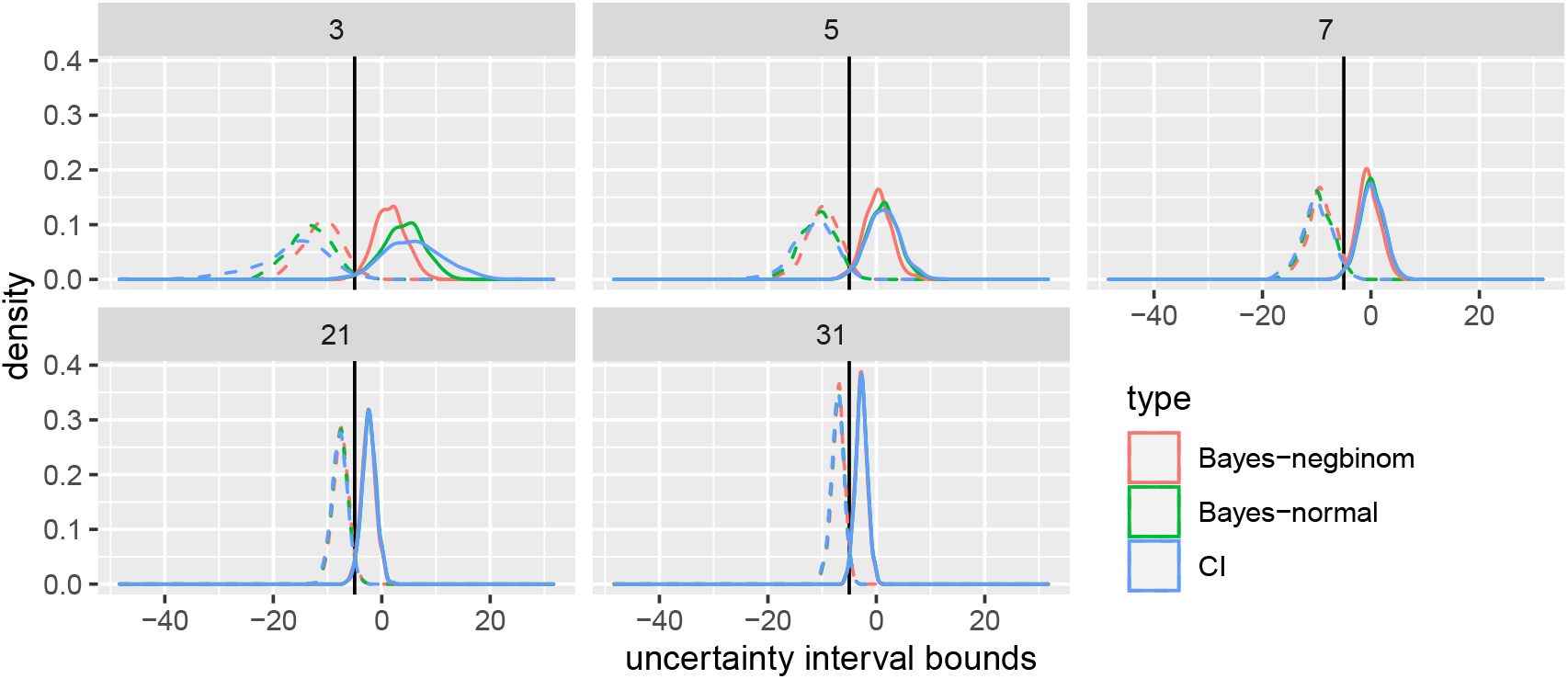
Distributions of lower (dashed lines) and upper (solid lines) bounds of uncertainty intervals for effect size *δ*. Red: 95% HDI for negative binomial model; green: 95% HDI for normal model; blue: 95% CI. True effect size *δ* = −5 marked by vertical black line. Numbers in top stripes are sample sizes *n* per mouse strain. Only data for the five sample sizes are shown, *n* = 3, 5, 7, 21, 31.

For the two smallest sample sizes, *n* = 3 and *n* = 5, the distributions for the upper bounds have most of their density in the positive, which means that we cannot exclude zero or positive effect sizes. This changes for the higher *n* (lower row in Fig. 4), for which positive effects become very unlikely, so that, if we were just interested in the *sign* of the effect, the highest *n* would not be needed.

Figure 5 compares the uncertainty interval widths for the three methods at all tested sample sizes *n*. For all *n*, the 95% HDIs from the negative binomial models are, on average, the shortest of the three uncertainty intervals and also have the least variation in lengths. The 95% CI and 95% HDI of the normal model are higher and more similar, especially for higher *n*, and in general vary more strongly. This means that if we are interested in minimizing uncertainty, as measured by the length of the uncertainty interval, we should prefer the more accurate negative binomial model for all *n*.

**Figure 5:**
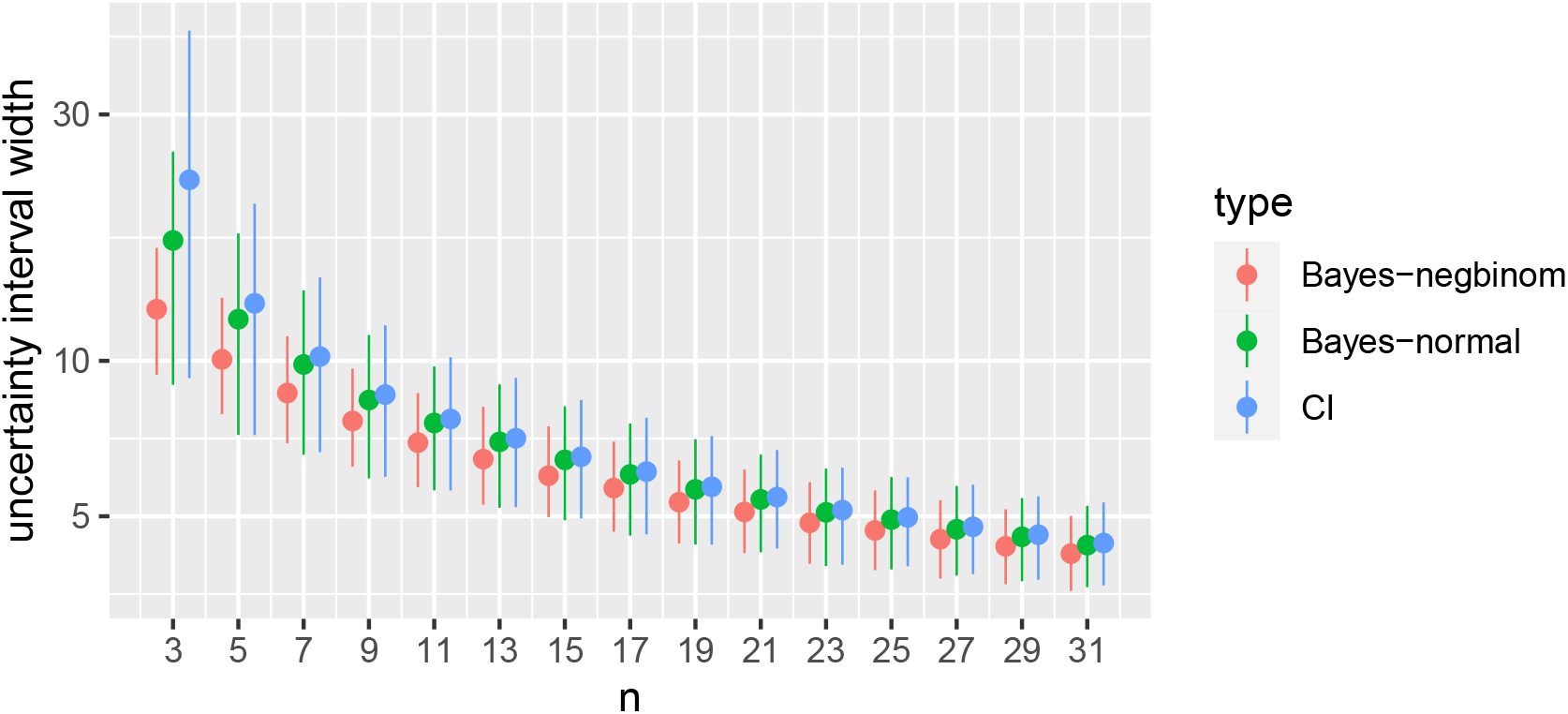
Comparison of uncertainty interval widths: 95% CIs (blue), 95% HDI for normal (green) and negative binomial (red) models for simulated samples of *n = 3, 5, 7,..*., 31 animals for each of the two strains A and B. Vertical axis is log-scaled. Bars cover 0.05 to 0.95 quantiles of interval lengths, dots are mean interval lengths.

### 6.2. Better model accuracy leads to more true positives

A short uncertainty interval is only valuable if it actually contains the true effect size. Hence, one crucial aspect of information gain yet to be quantified is whether an uncertainty interval contains the true effect size (here: *δ* = −5), but not zero. If it does, we call this a true positive (TP). If it does not, we call this a false negative (FN). As in our case we are sure, by construction of our computational experiment, that there is a true non-zero effect, there are no false positives or true negatives. In the following we focus on the false negative rate “fnr”, i.e. the fraction of FNs among all the 1000 A vs B tests for an *n*. Because FNs and TPs are complementary, the true positive rate is just (1 – fnr). From the data in our computational experiments described above we can compute one fnr value for each *n*, the number of tests *i* for that *n* that lead to a FN divided by 1000, the total number of tests. To assess the reliability of this value, we carry out for each *n* 100 resamplings of 1000 FNs and TPs and compute the fnr value for each of these bootstraps. This gives us for each *n* a distribution of fnr values that we characterize with its 0.05, 0.5, and 0.95 quantiles (Fig. 6).

**Figure 6:**
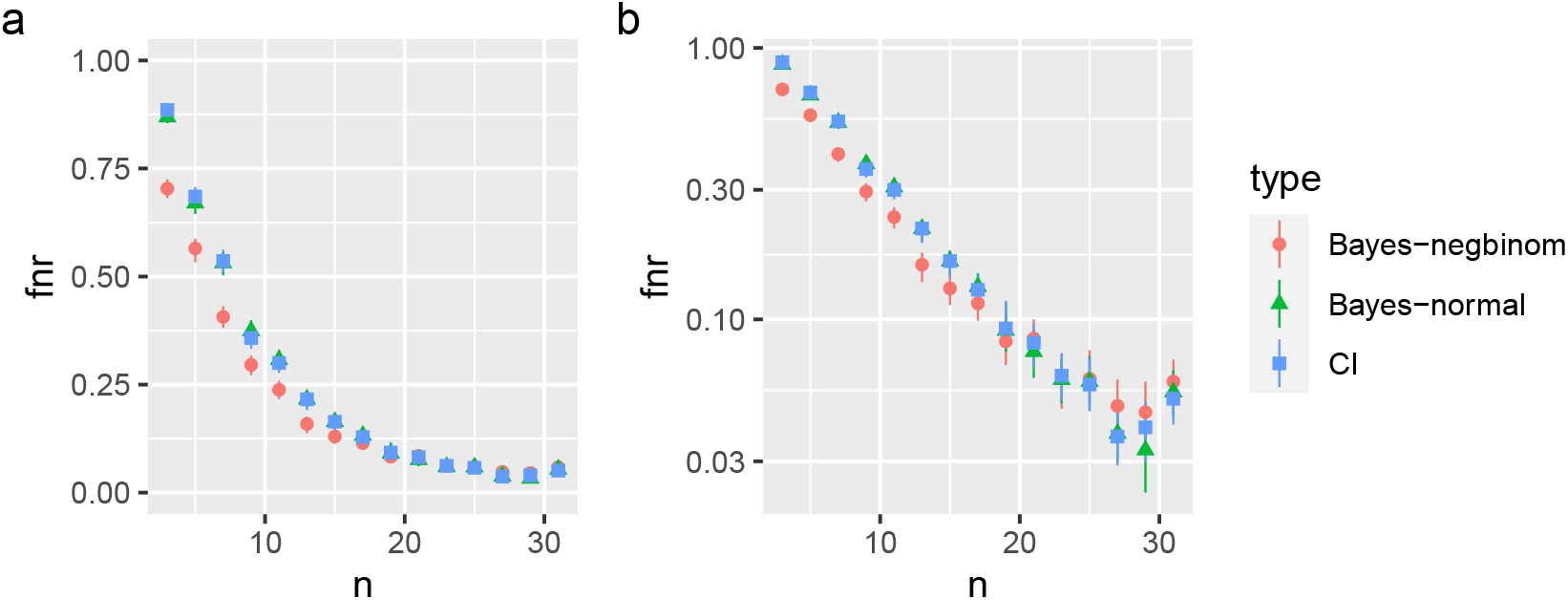
False negative rate (fnr) for each of the sample sizes *n* = 3, 5,…, 31 per strain, and method: 95% HDI of negative binomial models (red), 95% HDI for normal models (green), 95% CI (blue). (a) linear fnr axis, (b) logarithmic fnr axis. Dots are median fnr values of 100 bootstraps, bars reach form 0.05 to 0.95 quantiles of bootstrapped fnr values.

The effect of the model accuracy is substantial with the negative binomial model having the lowest fnr values, especially for *n* ≤ 15. Thus, this model has not only the most narrow uncertainty intervals, as we have seen in Fig. 5, but also most often encloses the true value of *δ* = −5 in these uncertainty intervals. There is little fnr difference between the 95% CIs and 95% HDIs of the normal model, possibly because both rely on a normal model of the data.

In the log-scaled Fig. 6b we see that, up to about *n* = 15, the fnrs for all methods decay exponentially with increasing *n*, with a constant ratio of 0.8 between the fnr of the 95% HDIs of the negative binomial model and the other two methods, i.e. the fnr is consistently 20% lower for the more accurate negative binomial model.

For larger *n* we reach FNR levels in the single digit percent range (Fig. 6). At these levels, FNR differences between the different models and methods are dominated by the noise from the limited sampling in our computer experiment, and uncertainty intervals between models and methods are no longer separated.

In terms of mouse number reduction, we see in Figs. 5 and 6 that switching from the less accurate normal model to the more accurate negative binomial model corresponds to a shift of *n* by about two, i.e. we save about two animals per strain or four animals per experiment. This demonstrates again that with the more accurate model, we gain more information with the same number of animals, or gain the same information with less animals.

Figure 6 or the corresponding tabulated values can be used for planning an experiment with the minimum number of animals that allows detection of the effect with a given probability (“power”). For instance, under the assumption of an effect of size *δ* = −5 between the negative binomial distributions in groups A and B, we want to determine the minimum number *n* of animals that allow detection of this effect with a power of 0.8, equivalent to fnr = 0.2. We look up in Fig. 6 the smallest *n* for which fnr ≤ 0.2, which is about *n* = 12 (interpolating between red points at *n* = 11 and *n* = 13). Thus, for the given effect size we would need 12 animals in each of the two groups A and B to detect the effect with a power of 0.8.

## 7. Outlook and conclusions: steps towards more accurate models in practice

For many common experiments in animal research, models with better accuracy, as exemplified in the present work, are already available in theory. This is not limited to the set of classical Bayesian models for which analytical posteriors are known, but it includes a much larger set of models that can be fitted computationally. If a model of the posterior probability can be mathematically formulated and implemented in probabilistic programming languages such as Jags [25] or Stan [22], Bayesian inference can in practice often be applied successfully. We have found this to be true for dozens of types of animal experiments commonly used in the literature. This includes experiments for which test assumptions of the usual NHSTs are approximately fulfilled, but also experiments that clearly violate typical NHST assumptions such as normality, continuity, homogeneity of variances, etc., as demonstrated with the example in this manuscript. In fact, Bayesian estimation supersedes the t-test even if test conditions are fulfilled [26].

The main limitation of Bayesian modeling that we encounter in practice is model complexity. Firstly, with increasing complexity, it becomes more and more difficult to write down and implement a scientifically plausible model. For the simpler type of model elements used in Bayesian networks, this problem can be solved efficiently by structure learning algorithms [27]. This is currently not possible for the more general approach taken in this manuscript. Secondly, efficient numerical Bayesian inference requires numerical Methods such as Hamiltonian Monte Carlo [28]. Such algorithms usually come with a set of mathematical requirements that may not be fulfilled, which can lead to inefficient or aborting sampling. Third, even if a model can be properly formulated and efficiently sampled, the posterior may be too large for a sufficiently thorough sampling. However, for most of the applications in the current animal research literature that are typically analyzed by NHSTs, these limitations are irrelevant as the applicable Bayesian models have relatively low complexity and can be sampled efficiently and thoroughly.

As we have demonstrated here, the application of more accurate models would improve information gain per animal and thus contribute to a reduction of animal numbers. Nevertheless, the most frequently used analysis in the literature is not Bayesian inference but NHST, such as the t-test used in this manuscript to compute confidence intervals.

In our experience, there are several reasons that may account for this dismal state: Research practitioners in the life sciences …

1. have been trained with NHSTs that are easy to use in a black-box recipe manner, and they stick to what they have learned;
2. have no clear understanding of NHST and model limitations;
3. appreciate the seeming clarity and certainty produced by NHST procedures with their classification of results into “significant” and “not significant”;
4. think that NHSTs are required for publication, and some reviewers indeed require NHSTs;
5. do not know alternative statistical methods;
6. do know alternative statistical methods but do not have access.

There are at least two lines of action to respond to these points: better education and better software. The first line needs great efforts and the time scale of a generation to succeed. But the vivid discussion in the scientific literature [15, 12] makes us hopeful. The new insights into experimental planning and evaluation will over time permeate all relevant aspects of education and scientific practice, from university programs to editorial boards and funding agencies. The second line, better software, means that advanced methods and models should become more accessible for science practitioners without special computational skills. Tools like our own BAYAS (https://bayas.zmb.uni-due.de) are under development to make this possible. We suspect that both lines of action will mutually reinforce as they advance: better education lays the groundwork for a conceptual understanding of better methods, and easier access to better methods will make education and practical application easier.

The Bayesian negative binomial model presented in this work has increased information gain per animal despite the relatively simple model structure. One of the advantages of the Bayesian approach that it facilitates formulation of more complex models. If we exploit this feature and map further aspects of the experimental system that contribute to the variation of *y* onto a Bayesian model, we will be able to estimate the contribution of the mouse strain to *y* even more accurately, i.e. increase information gain further. This is not a mere theoretical possibility: the experiment reported in [6] had a batch structure, i.e. groups of animals were in some respects more similar, that contributes to the overall variation. Accounting for that batch structure increases model accuracy, as will be detailed elsewhere.

Another dial that can be turned in Bayesian inference to increase information gain is the prior. The prior in our model had been *weakly informative*. However, if we have specific information from the literature, or if we increase the number of animals batch by batch (“Bayesian updating”), we can provide much narrower *informative priors*, leading to efficient information accumulation, as we will demonstrate in work that is underway.

## Acknowledgments

CW and DH gratefully acknowledge funding by the Federal Institute for Risk Assessment (BfR) with grant number 60-0102-01.P584.

## Appendix A: normal model

The Stan program of the normal model:

~~~
**data** {
  **int**<**lower** =0> n; *// table rows (animals*)
  **int** < **lower** =0> s[n]; *// 2 conditions (e.g. strains*)
  **real** y[n ] ; *// number of observations of behavior*
}
**parameters** {
  **vector** [2] mu; *// one mu per condition*
  **real** < **lower**=0> sig ; *// sd*
}
**model** {
  mu ~ **normal** ( 0 . 0 , 1 0 ) ;
  sig ~ **exponential** (0.1);
  **for** ( i **in** 1 : n) {
    y[i] **~ normal**(mu[s[i]], sig);
  }
}
**generated quantities** {
  **real** yrep [n ] ;
  **real** lo g_li k [n ] ;
  **for** ( i **in** 1 : n) {
    yrep[i] = **normal_rng** (mu[s[i]], sig);
    log_lik[i] = **normal_lpdf**(y[i] | mu[s[i]], sig);
  }
}
~~~

## Appendix B: negative binomial model

The Stan program of the negative binomial model, formulated as a generalized linear model with an exponential inverse link function:

~~~
**data** {
  **int** < **lower** = 1> n;
  **int** <**lower** =0> y[n]; *// center entries*
  **int** < **lower** =0> s[n]; *// strain, 1 for A, 2 for B*
}
**parameters** {
  **real** < **lower**=0> phi;
  **real** alpha [2];
}
**transformed parameters** {
  **real** < **lower**=0> mu[2];
  **for** ( i **in** 1 : 2) {
    mu[i] = **exp**(alpha[i]);
  }
}
**model** {
  alpha **~ normal**(0 , 2) ;
  phi **~ exponential** (0.05);
  **for** ( i **in** 1 : n) {
    y[i] **~ neg_binomial_2** (mu[s[i]], phi);
  }
}
**generated quantities** {
  **int** yrep [n ] ;
  **real** lo g_li k [n ] ;
  **for** ( i **in** 1 : n) {
    yrep[i] = **neg_binomial_2_rng** (mu[s[i]], phi);
    log_lik[i] = **neg_binomial_2_lpmf**(y[i] | mu[s[i]], phi);
  }
}
~~~

## Footnotes

1 We use the definition of skewness in [7], i.e. 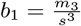, with *m*_3_ the third moment after centering and *s* the standard deviation.

